# Virus-driven tRNA competition universally represses host genes with similar codon usage

**DOI:** 10.64898/2026.07.02.736060

**Authors:** Feng Chen

## Abstract

Viral infection triggers a competition for transfer RNAs between viral and host endogenous genes, often leading to the repression of endogenous gene translation. However, how endogenous genes are affected by this competition remains unclear. Three possible answers are considered: the abundant-tRNA shortage hypothesis, the rare-tRNA shortage hypothesis, and the viral similarity repression hypothesis. Using pan-virus Ribo-seq data, it is found that endogenous genes whose codon usage bias (CUB) matches host tRNA supply or viral CUB are strongly repressed due to a positive correlation between endogenous CUB-tRNA mismatch and endogenous-viral CUB difference, supporting the abundant-tRNA shortage and the viral similarity repression hypotheses. In *E. coli* manipulative experiments with synonymous gentamicin resistance proteins, this positive correlation supports the abundant-tRNA shortage hypothesis, a non-positive correlation supports the rare-tRNA shortage hypothesis, and both positive and non-positive correlation support the viral similarity repression hypothesis. Finally, analysis of human virus genomes reveals this positive correlation for most viruses, but a non-positive correlation is found in a few, reflecting the diversity of virus–host interaction strategies. These findings establish viral similarity repression as a universal principle, a previously unrecognized complexity in the diverse coevolution of virus and host.

## Introduction

During infection by animal viruses (excluding bacterial viruses that encode their own tRNAs ^1,2^), the virus hijacks the host’s extremely scarce translational resources ^3,4^ to massively produce viral proteins, creating intense competition for tRNAs between viral and endogenous genes ^5,6^. Due to the differential tRNA supply in the cell, synonymous codons are not used with equal frequencies, a phenomenon termed codon usage bias (CUB) ^7–9^. The influence of CUB on translation efficiency through tRNA competition can be divided into cis-regulation and trans-regulation. Cis-regulation refers to the effect of CUB on the translation of the viral gene itself, with a smaller CUB difference between the viral gene and the host enhancing its own translation ^10–12^. Trans-regulation refers to the effect of CUB on the translation of endogenous genes, where a high similarity between the viral gene’s CUB and that of the host depletes host tRNAs and consequently represses the translation of endogenous genes ^13–15^. Together, cis-regulation determines how many viral protein molecules are produced, while trans-regulation determines how strongly the tRNA consumption during the synthesis of each viral protein molecule represses endogenous translation. This trade-off between cis- and trans- regulation of CUB ^16^ becomes particularly acute during viral infection, raising a critical question: which type of endogenous genes suffer the strongest translational repression for a given virus infection, and why?

To address this question, three potential hypotheses were proposed to explain how viral translation differentially inhibits endogenous genes by selectively depleting the tRNA pool. First, the Rare tRNA Shortage (RTS) hypothesis posits that massive viral translation primarily depletes rare tRNAs due to their low basal abundance ^17^, rendering them even scarcer. Under this scenario, endogenous genes that rely on rare tRNAs would experience disproportionately stronger inhibition (Fig. 1a). Second, the abundant-tRNA shortage (ATS) hypothesis considers the evolutionary scenario where viral codon usage converges with that of the host ^12,18,19^, leading to preferential consumption of abundant tRNAs ^20,21^. Under this scenario, endogenous genes that rely on abundant tRNAs would experience disproportionately stronger inhibition (Fig. 1b). Third, the viral similarity repression (VSR) hypothesis posits that the repression strength of an endogenous gene is determined by the overall codon usage similarity between the gene and the virus, regardless of whether the tRNAs involved are rare or abundant. Specifically, genes whose codon usage is most similar to that of the virus (i.e., with a small CUB distance to the virus) are predicted to be most strongly repressed, because their tRNA demand overlaps most with that of the virus (Fig. 1c).

**Fig. 1.**
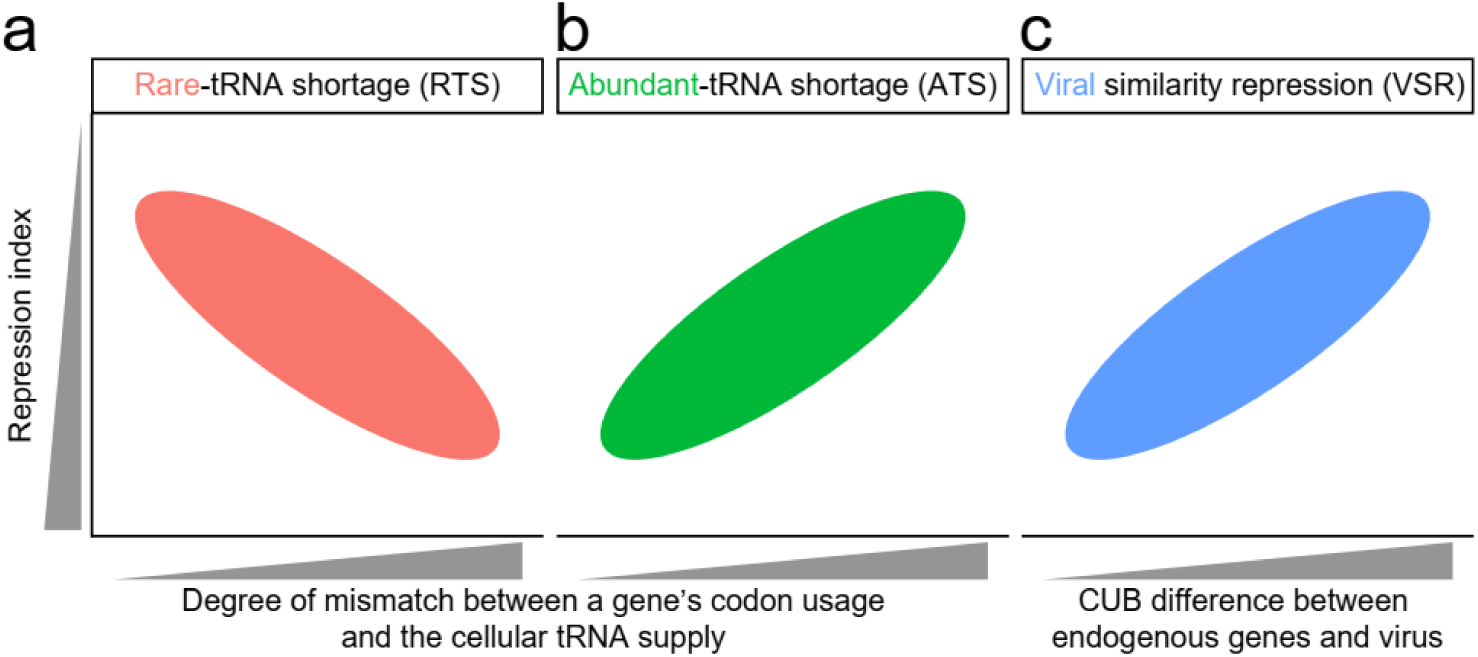
Three competing hypotheses for viral translational repression on endogenous genes. (a) Rare-tRNA shortage (RTS). Upon viral infection, the supply of rare tRNAs is drastically reduced, while abundant tRNAs remain relatively sufficient. Endogenous genes that rely on rare tRNAs (i.e., large values of *x*-axis) are therefore more strongly translationally repressed. (b) Abundant-tRNA shortage (ATS). Under evolutionary pressure, viral codon usage converges to host-abundant codons, making the virus highly dependent on abundant tRNAs. Infection leads to severe depletion of these abundant tRNAs due to massive viral translation. Consequently, endogenous genes that use abundant tRNAs (i.e., small values of *x*-axis) experience stronger repression. (c) Viral similarity repression (VSR). Upon infection, both rare and abundant tRNAs become depleted, but to different extents depending on viral sequence characteristics (e.g., tRNA demand). Endogenous genes whose codon usage is most similar to that of the virus (i.e., small values of *x*-axis) are most strongly repressed, because their tRNA demand overlaps most with that of the virus. In panels (a) and (b), the *x*-axis represents *D*, a measure of the disparity between a gene’s codon usage and the cellular tRNA supply (see Methods), and the *y*-axis represents the extent of viral translational repression index on endogenous genes (lower values indicate stronger repression, see methods). In panel (c), the *x*-axis represents *V*, a measure of the CUB difference between endogenous genes and viruses (see Methods), while the *y*-axis is the same as in panels (a) and (b).

To test these hypotheses, Ribo-seq data from human A549 cells transfected with a plasmid library of viral genes were analyzed. Endogenous genes whose codon usage matches the host tRNA supply or the viral codon usage were found to be strongly repressed, a result that simultaneously supports the ATS and VSR hypotheses. A mechanistic dissection further revealed that the collective tRNA demand of the viral library approximates the host tRNA supply, driving a strong positive correlation between endogenous-viral CUB difference (denoted as *V*) and endogenous CUB-tRNA mismatch (denoted as *D*). Manipulative experiments using *E. coli* strains expressing four synonymous gentamicin resistance proteins (*GmR*) further supported the pan-viral findings: when the correlation between endogenous *V* and *D* was positive, endogenous genes with small *V* or small *D* were more strongly repressed (supporting VSR and ATS); when the correlation was non-positive, endogenous genes with small *V* and large *D* were more strongly repressed (supporting VSR and RTS). Extending this framework to real human viral genomes, a positive *V*-*D* correlation for endogenous genes was found in most viruses, while a negative *V*-*D* correlation for endogenous genes was found in five viruses, indicating that virus–host interaction strategies are diverse. Collectively, these results establish VSR as a universal principle that provides a unifying explanation for the diversity of viral-host coevolution.

## Results

### Correlation between codon usage bias and translational repression index of each endogenous gene

To discriminate among the three hypotheses (Rare-tRNA shortage, RTS; Abundant-tRNA shortage, ATS; Viral similarity repression, VSR), Ribo-seq data from A549 cells ^22^ (seven repeat samples; see Table S1) transfected with a plasmid library of pan-viral genes (see Table S2) were reanalyzed. Repression index (lower values indicate stronger repression; see Fig. 1, *y*-axis and methods) of each endogenous gene due to viral gene translation was quantified as the fold change of transcripts per million of ribosome-protected fragments in transfected and wild type cells. For a series of translation efficiency percentile thresholds (from 100% of all genes down to the top 1%, based on TPM values of RPF; see methods), Spearman correlation coefficients were calculated between repression index and three CUB metrics, for which each endogenous gene has a value: *D*_opt_ (a metric of CUB-tRNA mismatch estimated from highly expressed genes; see methods), *D*_tRNA_ (a metric of CUB-tRNA mismatch estimated from tRNA expression; see methods), and *V* (a metric of CUB distance to virus; see methods).

Due to the limited throughput of Ribo-seq technology, the measurement of translation efficiency for genes with low translational activity may be inaccurate. Consequently, as translational activity increases, the observed pattern becomes more reliable. It was observed that with increasing translational activity (Fig. 2a, *x*-axis), the positive correlation between repression index and *D_opt_* of endogenous genes becomes stronger (Fig. 2a, *y*-axis). This trend cannot be attributed to a reduction in gene number, as random downsampling to the same number of genes does not reproduce the same observation (Fig. 2a, gray bars). The same finding was confirmed in other biological repeats of this dataset (Fig. S1, left panel). Using an alternative metric of CUB-tRNA mismatch (*D_tRNA_*), the consistent results were obtained (Fig. 2b. The other repeats in Fig. S1, middle panel). Together, these results support the notion that upon viral gene expression, highly abundant tRNAs become scarcer, leading to stronger translational repression of endogenous genes that have a smaller CUB-tRNA mismatch, consistent with ATS (abundant-tRNA shortage; Fig. 1b).

**Fig. 2.**
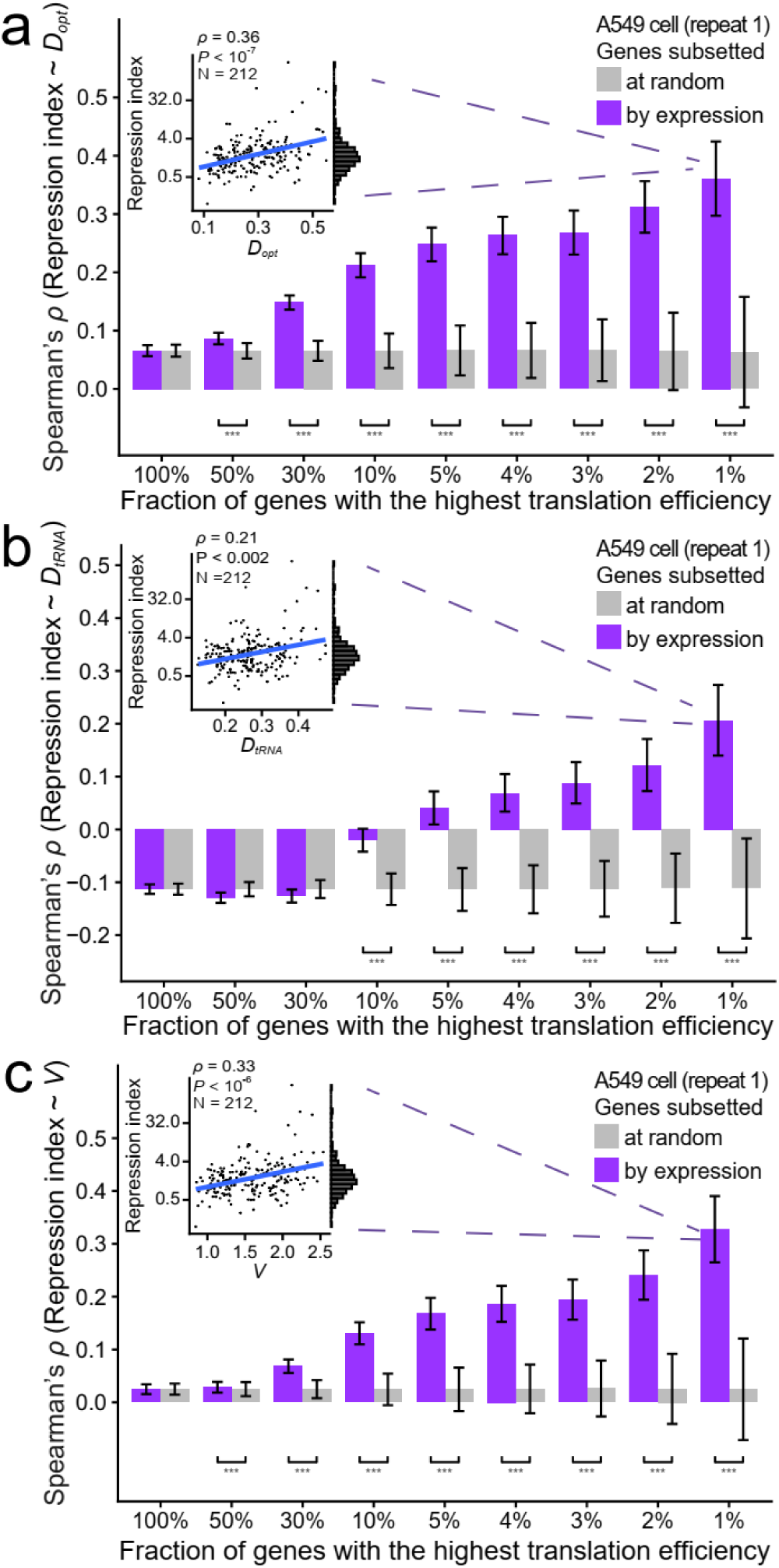
Pan-virus Ribo-seq data support hypothesis 2 (ATS) and hypothesis 3 (VSR). (a–c) For endogenous genes whose translation efficiency (TPM of RPF counts) exceeds a given threshold (*x*-axis), the Spearman’s rank correlation coefficient (y-axis) was estimated between the degree of viral translational repression index (see Methods) on endogenous genes and each of three CUB metrics: *D_opt_* (a), *D_tRNA_* (b), and *V* (c). Purple bars indicate results from the real gene set above the threshold, while gray bars represent results from a randomly sampled gene set of the same size as the real set. Error bars indicate standard deviations of the correlation assessed from 1,000 bootstraps (purple) or random samples (gray). The inset scatter plot in each panel shows one representative correlation, and the histogram to the right shows the distribution of repression index across genes. Statistical significance from a one-tailed Wilcoxon signed-rank test comparing real and random gene sets is indicated as *: *P* < 0.05, **: *P* < 0.01, or ***: *P* < 0.001.

It was further observed that with increasing translational activity (Fig. 2c, *x*-axis), the positive correlation between repression index and *V* of endogenous genes becomes stronger (Fig. 2c, *y*-axis), indicating that endogenous genes with higher CUB similarity to the virus are more strongly translationally repressed. This trend cannot be attributed to a reduction in gene number, as random downsampling to the same number of genes does not reproduce the same result (Fig. 2c, gray bars). The same finding was confirmed in other biological repeats of this dataset (Fig. S1, right panel). Together, these results support the notion that upon viral gene expression, tRNAs consumed by the viral genes become scarcer, leading to stronger translational repression of endogenous genes whose CUB profile more closely matches that of the virus, consistent with VSR (Viral similarity repression; Fig. 1c).

In summary, the pan-virus Ribo-seq data provide evidence supporting ATS and VSR, whereas no evidence supporting RTS (Rare-tRNA shortage) was found in the current dataset.

### The tRNA demand of pan-viral genes approximates host tRNA supply

Using pan-virus Ribo-seq data, evidence supporting both ATS (abundant-tRNA shortage) and VSR (viral similarity repression) was found. Despite their different mechanisms, why did the observed data support both? To explore this, the CUB characteristics and translation efficiency of pan-viral genes were further analyzed.

The *D_opt_* values ranged from 0.058 to 0.684 (median 0.316) for viral genes and from 0.032 to 0.732 (median 0.218) for endogenous genes (Fig. 3a). The *D*_tRNA_ values ranged from 0.094 to 0.747 (median 0.369) for viral genes and from 0.076 to 0.738 (median 0.237) for endogenous genes (Fig. 3b). Incidentally, the CUB-tRNA mismatch is significantly larger for viral genes than for endogenous genes (Fig. 3a-3b, Wilcoxon test *P* < 10^−300^). This observation does not imply that the tRNA consumption of viral genes differs from that of the host, because it does not account for the large differences in the number of protein molecules produced by each viral gene due to varying translation efficiencies. To resolve this, the translation efficiency of each viral gene was next calculated (see method) and it was found that viral gene translation efficiency was significantly negatively correlated with its CUB-tRNA mismatch value (Fig. 3c; *D_opt_*, upper panel; *D*_tRNA_, lower panel). This indicates that viral genes with a small CUB-tRNA mismatch produce many more protein molecules, while those with a large CUB-tRNA mismatch produce few. By definition, during the synthesis of a single protein molecule, genes with a small CUB-tRNA mismatch (e.g., *D_opt_* or *D_tRNA_* values around 0.2) consume high-abundance tRNAs in relatively larger amounts per protein, whereas genes with a large CUB-tRNA mismatch (e.g., *D_opt_* or *D_tRNA_* values 0.6) consume rare tRNAs in relatively larger amounts per protein. Consequently, considering all protein molecules synthesized, the total consumption of high-abundance tRNAs is large (due to both high per-protein demand and high protein output), while the total consumption of rare tRNAs remains small (because the low protein output of large CUB-tRNA mismatch viral genes outweigh their high per-protein demand). Therefore, although the CUB-tRNA mismatch values of pan-viral genes are larger than that of endogenous genes (Fig. 3a-3b), the total tRNA consumption is still dominated by viral genes with a small CUB-tRNA mismatch. In other words, the tRNA demand of pan-viral genes approximates the host tRNA supply.

**Fig. 3.**
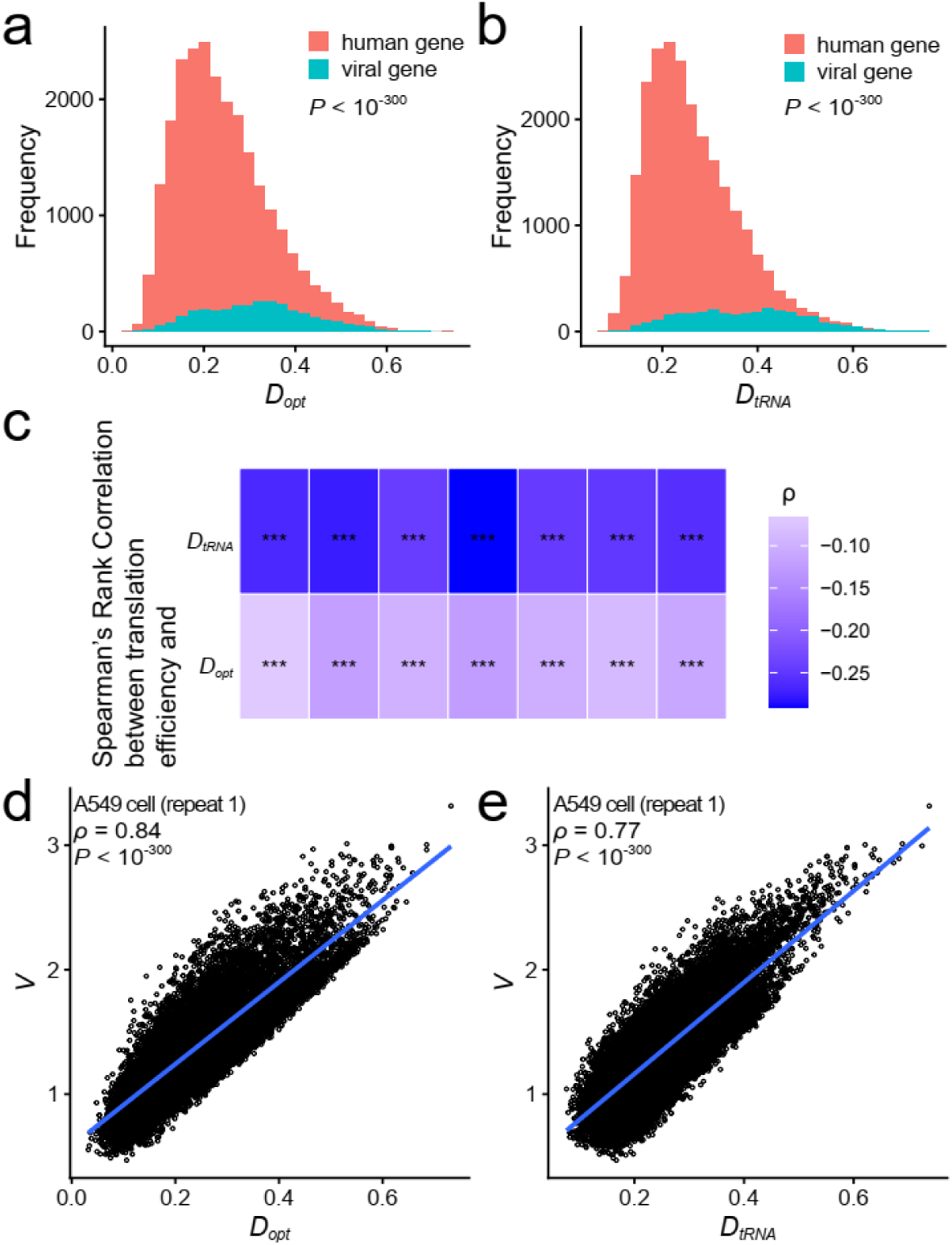
CUB-tRNA mismatch and translation efficiency of pan-viral genes. (a) Distribution of *D_opt_* values for pan-viral and endogenous genes. *D_opt_* is a metric of CUB-tRNA mismatch estimated from highly expressed genes (see Methods). (b) Distribution of *D_tRNA_* values for pan-viral and endogenous genes. *D_tRNA_* is a metric of CUB-tRNA mismatch estimated from tRNA expression (see Methods). For both panel (a) and (b), the Wilcoxon test *P* value were presented. (c) Spearman rank correlation between viral gene translation efficiency and CUB-tRNA mismatch measured by *D_opt_* (upper panel) or *D_tRNA_* (lower panel) in seven repeats (*x*-axis) of pan-virus Ribo-seq data. The negative correlation indicates that genes with smaller CUB-tRNA mismatch have higher translation efficiency. Statistical significance from Spearman rank correlation is indicated as ***: *P* < 0.001. (d) Spearman correlation between *V* (CUB distance to virus) and *D_opt_* for endogenous genes. (e) Spearman correlation between *V* and *D_tRNA_* for endogenous genes. For both panel (d) and (e), the Spearman’s rank correlation coefficient (ρ) and its two-tailed *P* value were presented. Each dot represents an endogenous gene, and the blue lines represent linear models fitted towards either *D_opt_* or *D_tRNA_*.

Based on the logic above, a given endogenous gene whose CUB is more similar to that of viral genes (i.e., a smaller *V* value) will also be more similar to the host tRNA supply (i.e., smaller *D_opt_* or *D*_tRNA_). To test this inference, Spearman correlation coefficients were calculated and *V* was found to be significantly positively correlated with both *D_opt_* (Fig. 3d; other repeats in Fig. S2a) and *D*_tRNA_ (Fig. 3e; other repeats in Fig. S2b). Thus, the CUB composition of pan-viral genes shapes their tRNA consumption to match host tRNA supply, explaining why both ATS and VSR are supported by the same dataset.

### CUB-tRNA mismatch of heterologous genes determines the translational repression pattern of endogenous genes

To validate the core conclusion above, positive and negative validation experiments were performed using exogenous genes to model virus–host interactions. For positive validation, Ribo-seq data from *E. coli* strains ^16^ expressing two synonymous *GmR* variants with small CUB-tRNA mismatch (*D_opt_* = 0.127 and 0.278, see methods) were reanalyzed, expecting *V* to be significantly positively correlated with *D_opt_*/*D_tRNA_* (Fig.4a) and thus the repression pattern to support both VSR (viral similarity repression) and ATS (abundant-tRNA shortage). For negative validation, Ribo-seq data from the same dataset ^16^, using two strains whose *GmR* variants have large CUB-tRNA mismatch (*D_opt_* = 0.778 and 0.878, see methods) were reanalyzed, expecting *V* not to be significantly positively correlated with *D_opt_*/*D_tRNA_* (or even negatively correlated, Fig. 4b) and thus the repression pattern to support both VSR and RTS (rare-tRNA shortage).

**Fig. 4.**
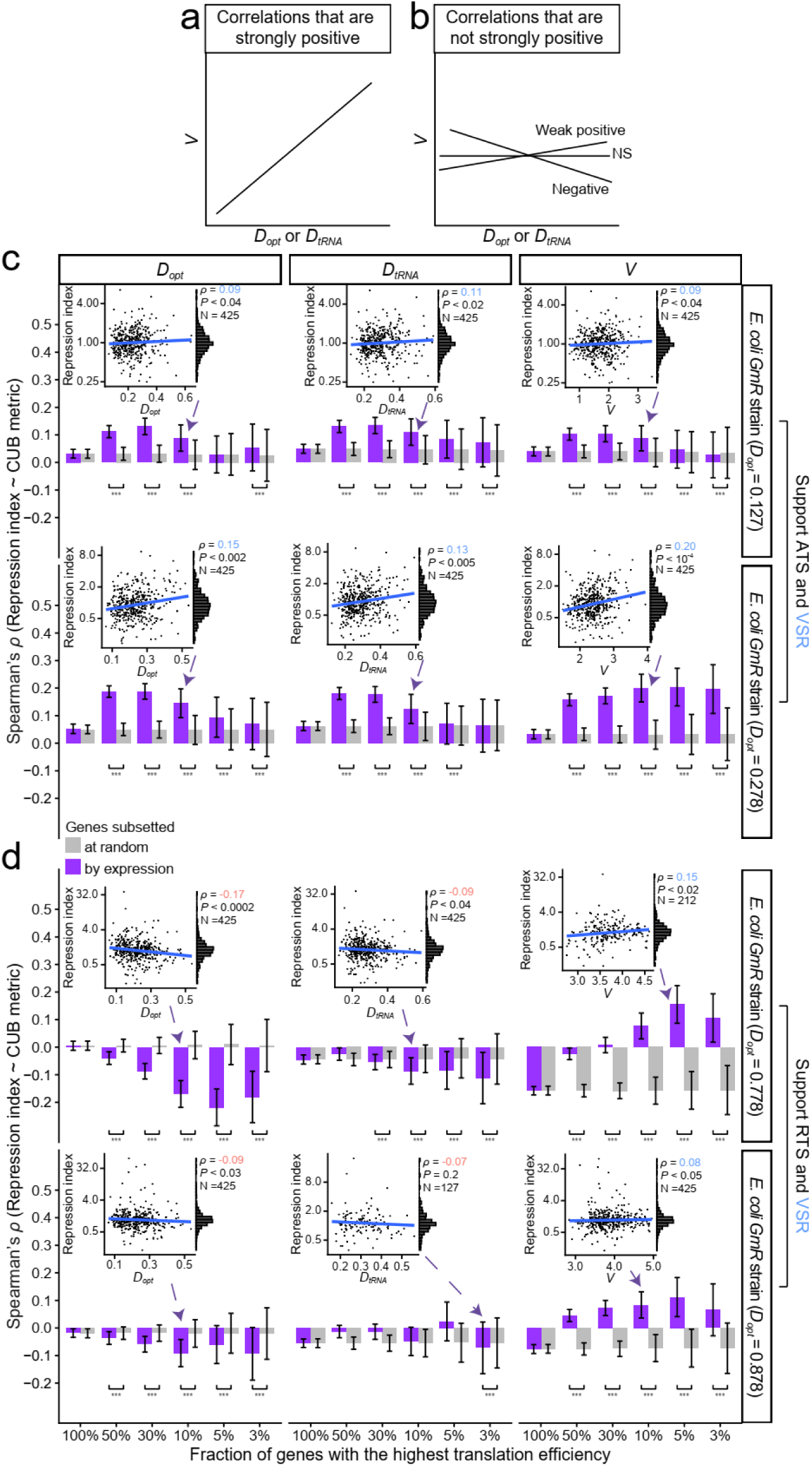
Validation of the mechanism linking exogenous gene CUB-tRNA mismatch to endogenous gene repression patterns. (a) Schematic of positive validation: an exogenous gene with small CUB-tRNA mismatch is expected to yield a strong positive correlation between *V* and *D_opt_*/*D_tRNA_*, based on the mechanism established in Fig. 3. (b) Schematic of negative validation: an exogenous gene with large CUB-tRNA mismatch is expected to yield no positive correlation (or negative correlation) between *V* and *D_opt_*/*D_tRNA_*. (c) For *GmR* strains with small CUB-tRNA mismatch, Spearman correlations between repression index and *D_opt_* (left), *D_tRNA_* (middle), and *V* (right) were shown. (d) For *GmR* strains with large CUB-tRNA mismatch, one-tailed Spearman correlations between repression index and *D_opt_* (left), *D_tRNA_* (middle), and *V* (right) were shown. For both panels (c) and (d), purple bars indicate results from the real gene set above the threshold, while gray bars represent results from a randomly sampled gene set of the same size as the real set. Error bars indicate standard deviations of the correlation assessed from 1,000 bootstraps (purple) or random samples (gray). The inset scatter plot in each panel shows one representative correlation, and the histogram to the right shows the distribution of repression index across genes. To make the conveyed information more intuitive, positive and negative correlation coefficients in inset scatter plots are marked in green and red, respectively. Statistical significance from a one-tailed Wilcoxon signed-rank test comparing real and random gene sets is indicated as *: *P* < 0.05, **: *P* < 0.01, or ***: *P* < 0.001.

For the two *GmR* strains with small CUB-tRNA mismatch (*D_opt_* = 0.127 and 0.278), *V* (CUB distance between endogenous genes and *GmR*) and CUB-tRNA mismatch (*D_opt_* and *D_tRNA_*) were calculated for each endogenous gene. Spearman rank correlation analysis revealed that *V* was significantly positively correlated with both *D_opt_* and *D_tRNA_* (Fig. S3 a-b; ρ > 0.68, *P* < 10^−300^). The repression index was found to be positively correlated with *D_opt_* (Fig. 4c, left panels) and *D_tRNA_* (Fig. 4c, middle panels), supporting ATS, and also positively correlated with *V* (Fig. 4c, right panels), supporting VSR. Thus, the positive validation confirms the mechanism observed in the pan-viral dataset.

For the two *GmR* strains with large CUB-tRNA mismatch (*D_opt_* = 0.778 and 0.878), the correlations between *V* and CUB-tRNA mismatch (*D_opt_* and *D_tRNA_*) for endogenous genes were examined. Spearman rank correlation analysis showed that *V* was significantly negatively correlated with *D_opt_* (Fig. S3 c-d, left panel; ρ < −0.15, *P* < 10^−22^), and only weakly correlated with *D_tRNA_* (Fig. S3 c-d, right panel; ρ = 0.11, *P* < 10^−22^; ρ = 0.03, *P* = 0.05). The repression index was found to be negatively correlated with *D_opt_* (Fig. 4d, left panel) and *D_tRNA_* (Fig. 4d, middle panel). This negative correlation suggests that translation of the exogenous gene leads to increased scarcity of rare tRNAs, thereby preferentially repressing endogenous genes with larger CUB-tRNA mismatch (i.e., higher *D_opt_*/*D_tRNA_*), which is consistent with RTS (rare-tRNA shortage). Meanwhile, repression index was positively correlated with *V* (Fig. 4d, right panel), suggesting that tRNAs consumed by the exogenous gene become scarcer, leading to stronger translational repression of endogenous genes whose CUB profile more closely matches that of the exogenous gene, which is consistent with VSR (viral similarity repression). The Spearman correlation coefficients for repression index observed here (Fig. 4c-d) are smaller than those in the pan-viral data. For example, the maximal positive correlation between repression index and *D_opt_* across all translation efficiency thresholds was ρ = 0.36 in the pan-viral dataset (Fig. 2a; 1% genes with highest translation efficiency), compared to ρ = 0.19 in the *GmR* strains (Fig. 4c, second row, left panel; 30% genes with highest translation efficiency). This reduction can be attributed to the fact that only a single exogenous gene (*GmR*) is expressed in this system, leading to a much smaller consumption of translational resources compared to the pan-viral gene library. Thus, the negative validation confirms the mechanism observed: when the exogenous gene’s tRNA demand does not match the host tRNA supply, the repression pattern shifts to RTS and VSR.

Taken together, these positive and negative validations not only confirm the mechanism inferred from the pan-viral data, but also demonstrate that VSR is a universal principle, as it consistently holds regardless of the CUB-tRNA mismatch between the exogenous gene and the host. In contrast, ATS is supported only when the exogenous gene’s CUB-tRNA mismatch is small (resulting in a strong positive correlation between *V* and *D_opt_*/*D_tRNA_*), whereas RTS is supported only when the CUB-tRNA mismatch is large (where *V* is no longer positively correlated with *D_opt_*/*D_tRNA_*).

### Extending the framework to human virus

The validation experiments above demonstrate that the correlation between *V* and *D_opt_*/*D_tRNA_* is a key determinant of whether the repression pattern follows ATS/VSR (positive correlation) or RTS/VSR (non-positive correlation). To extend this framework, the following question was asked: among real human viruses, how often does a viral genome’s tRNA demand approximate the host supply, and how often does it deviate? Critically, in natural infections, the host is exposed to the entire viral genome rather than individual viral genes. Therefore, complete genome sequences of human viruses were downloaded from NCBI Virus (see methods). For each viral genome, *V* (a metric of CUB distance to virus; see methods), as well as the CUB-tRNA mismatch (*D*_opt_ and *D*_tRNA_; see methods) were calculated for each endogenous gene. The Spearman correlation between *V* and *D*_opt_/*D*_tRNA_ was then assessed. Through this correlation, the importance of distinguishing among the three hypotheses in the co-evolution of virus and human can be defined.

Consistent with the validation experiments above, it was observed that the direction of the correlation between endogenous *V* and *D_opt_* is negatively correlated with the *D_opt_* value of the viral genome itself (Fig. 5a; Spearman rank correlation coefficient ρ = −0.7, *P* < 10^−300^). Specifically, when the viral *D_opt_* is small (Fig. 5a, left side of *x*-axis), endogenous *D_opt_* and *V* are positively correlated (Fig. 5a, red points). Conversely, when the viral *D_opt_* is large (Fig. 5a, right side of *x*-axis), endogenous *D_opt_* and *V* tend to be negatively correlated (Fig. 5a, blue points). The same pattern was also observed using the alternative CUB-tRNA mismatch metric *D_tRNA_* (Fig. S4a, Spearman rank correlation coefficient ρ = −0.96, *P* < 10^−300^). The examples of negative correlation of Fig. 5a and Fig.S4a were shown in Fig. 5b and Fig.S4b, respectively. Among the 524 human virus species analyzed in this work, a very small fraction (5 out of 524; binomial test *P* < 10^−146^) exhibited a significant negative correlation between endogenous *D_opt_* and *V* (two representative examples of viruses are shown in Fig. 5c–d; the remaining viruses are listed in Table S3). Although these five viruses are few in number, they represent important human pathogens with serious clinical consequences. For example, rubella (*Rubivirus rubellae*) virus infection during pregnancy can lead to spontaneous abortion, stillbirth, congenital rubella syndrome ^23,24^. Neonatal HSV2 (*Simplexvirus humanalpha2*) infection, often transmitted by asymptomatic mothers with primary infection near delivery (risk ∼50%), is difficult to detect early and has a mortality rate of ∼60% without treatment ^25^. Using the alternative CUB-tRNA mismatch metric *D_tRNA_*, also a significant minority (140 out of 524; binomial test *P* < 10^−26^) of viruses exhibited a significant negative correlation between endogenous *D_tRNA_* and *V* (Fig. S4; viruses listed in Table S4), including well-known human pathogens such as monkeypox (*Orthopoxvirus monkeypox*) ^26^, SARS-CoV-2 (*Severe acute respiratory syndrome coronavirus 2*) ^27^, and HPV (*Human papillomavirus*) ^28^. Although the two metrics yield different absolute numbers due to the fact that *D_opt_* and *D_tRNA_* each deviate from the actual tRNA pool in different ways (see Methods), both indicate that the vast majority of viruses likely follow the ATS and VSR patterns, while only a small fraction may follow RTS and VSR. This distinction in virus–host translational interaction mechanisms highlight the importance of distinguishing among the three hypotheses for diverse virus-host co-evolution.

**Fig. 5.**
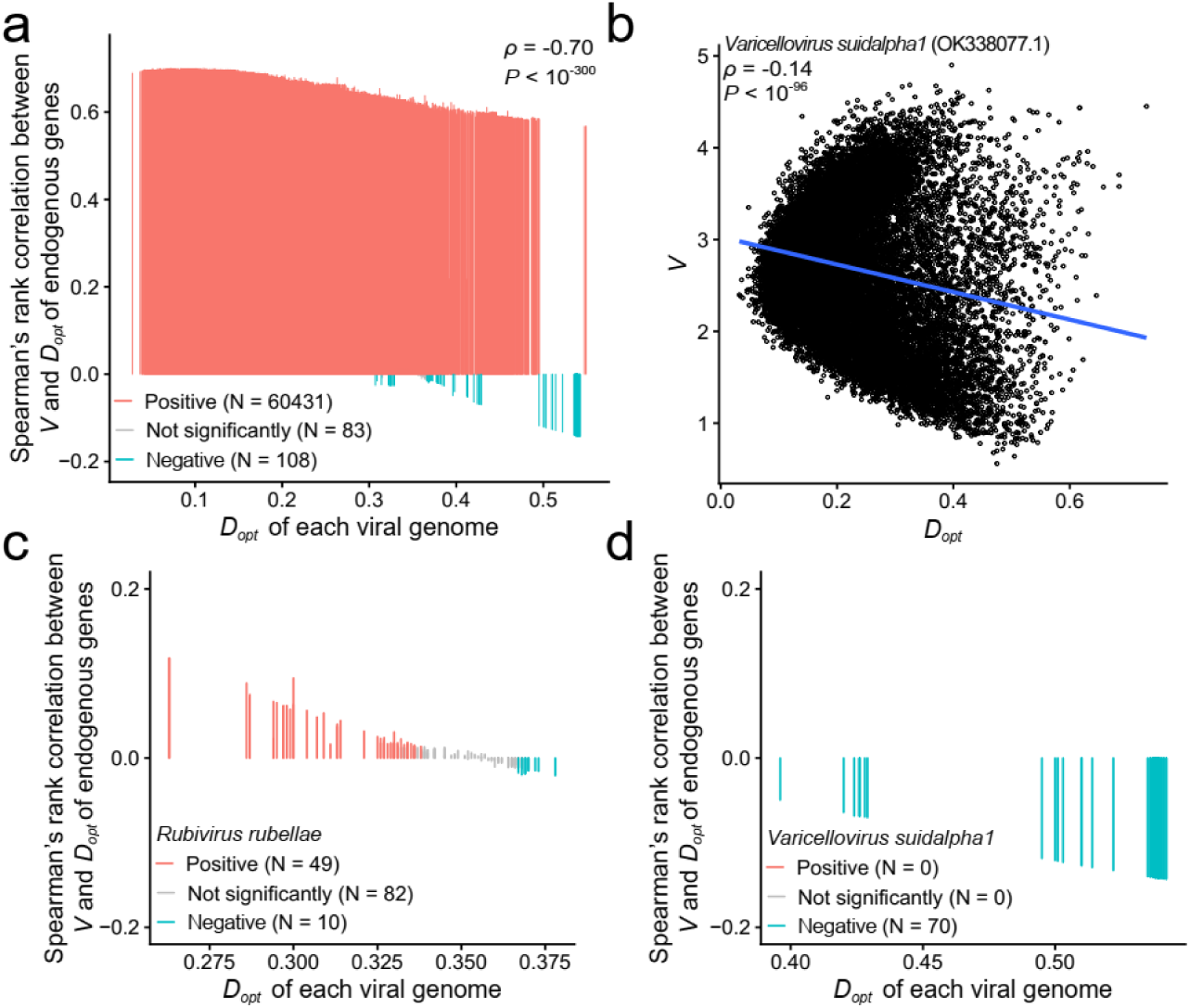
Viral *D_opt_* determines the direction of endogenous *V*–*D_opt_* correlation. (a) Lollipop chart showing the relationship between the *D_opt_* value of each human viral genome (*x*-axis) and the Spearman rank correlation coefficient between endogenous *V* and *D_opt_* (y-axis). Each vertical segment in the plot represents a single viral genome. (b) Representative example of a viral genome showing a negative correlation between endogenous *V* and *D_opt_*. The virus name and corresponding GenBank accession number are indicated in the panel. Each point represents an endogenous gene. For both (a) and (b), the Spearman’s rank correlation coefficient (ρ) and its two-tailed *P* value were presented. (c–d) Two representative viruses that exhibit a significant negative endogenous *V*-*D_opt_* correlation. Each vertical segment in the plot represents a single viral genome of the representative viruses.

## Discussion

In the current work, pan-virus Ribo-seq data, *E. coli* validation experiments, and genome data of human viruses were used to discriminate among three hypotheses of translation repression. The results consistently show that endogenous genes whose codon usage matches that of the virus are more strongly repressed, supporting the VSR hypothesis. The ATS hypothesis is supported when there is a positive correlation between *V* and *D* for endogenous genes under infection with a given human virus. In contrast, the RTS hypothesis is supported when the correlation is non-positive. Applying this framework to real human viruses, five viruses were found to display such negative correlation, and their neonatal severity (i.e., rubella and HSV-2) underscores the importance to distinguish among the three hypotheses.

Several limitations of our results merit discussion. First, pan-virus Ribo-seq data rather than data from a single virus were used. In this setting, tRNA depletion and endogenous gene repression represent the average effect of all viral genes combined, which helps to average out idiosyncratic regulatory mechanisms of individual viral genes and reveal core principles. However, because the plasmid library enters cells randomly, different cells may be dominated by different viral gene subsets, so the observed average repression index may deviate from the true effect in any given cell, potentially masking some patterns. This means that the actual repression effects are likely stronger than observed, and the fact that clear signals are still detected supports the robustness of the conclusions. Second, the experimental system used antibiotic resistance genes to model translation elongation, without considering factors such as translation initiation and immune responses that influence virus–host interactions. Therefore, the three hypotheses proposed here are primarily applicable to tRNA competition during translation elongation and do not negate other known interaction mechanisms identified in previous studies ^29^. Third, a notable discrepancy was observed between the two metrics of CUB-tRNA mismatch. Specifically, *D_opt_* identified five viruses whose infection will result in a negative *V*-*D* correlation of endogens gene, whereas *D_tRNA_* identified 140 such viruses. The Spearman correlation between viral *D_opt_* and the endogenous *V*-*D* correlation was –0.7 (Fig. 5a), whereas that using *D_tRNA_* was –0.96 (Fig. S4a). The non-overlapping 95% confidence intervals indicate a significant difference (Steiger’s *Z* test *P* < 10^−300^), suggesting that *D_tRNA_* more faithfully captures the biological pattern. Hence, the five viruses identified by *D_opt_* represent a conservative estimate, and the actual number of viruses following the RTS/VSR pattern is likely larger, as indicated by the *D_tRNA_* results (Table S4). Human viruses with significant clinical importance can be discovered by both metrics that support the RTS hypothesis, further underscoring the importance of distinguishing among the three scientific hypotheses.

A question worth discussing is why some viruses exhibit a codon usage that aligns with host abundant tRNAs (ATS), while others retain a mismatch that leads to rare-tRNA shortage (RTS). One possible explanation is that viral virulence is associated with the degree of CUB-tRNA mismatch ^15^. Specifically, viruses that preferentially use abundant tRNAs (small CUB-tRNA mismatch) tend to have high virulence, because they can rapidly produce large amounts of viral proteins and lead to tRNA depletion ^13^. In contrast, viruses that rely on rare tRNAs (large CUB-tRNA mismatch) tend to have low virulence, as their slower translation may limit replication speed. During virus–host co-evolution, highly virulent viruses may gradually accumulate mutations that increase their CUB-tRNA mismatch as a trade-off to reduce translational burden and host damage. A typical example is that the CUB of SARS-CoV-2 appears to be gradually dissimilating from the tRNA supply of human ^15,30^, which is consistent with reports that its virulence has decreased ^31,32^. Moreover, highly virulent viruses are more likely to cause severe symptoms and thus be sequenced and submitted, whereas low-virulence viruses often cause mild or asymptomatic infections and are under-represented. This sampling bias in public databases explains why the two metrics of CUB-tRNA mismatch (*D_opt_* and *D_tRNA_*) identify many more viruses supporting ATS (high virulence) than RTS (low virulence).

Our work has at least two important applications. First, it was found that the VSR hypothesis always holds, whereas whether ATS hypothesis or the RTS hypothesis is supported depends on whether the correlation between *V* and *D* is positive. This finding allows researchers to use metric *V* to predict the relative repression index of endogenous genes under viral infection, without the need for accurate assessment of the host tRNA pool which remains technically challenging (see Methods). For viruses with known genome sequences, the *V* metric can rapidly assess the relative susceptibility of different endogenous genes without the need for experimental measurement of the tRNA pool. Second, in the context of codon optimization for mRNA vaccines, traditional strategies often aim to perfectly match the vaccine gene’s codons with the host’s abundant tRNAs to maximize antigen protein expression. However, our previous work suggests that excessive CUB-tRNA matching may lead to overconsumption of tRNAs ^13,16^, potentially causing an unintended translational burden, which could help explain some of the adverse effects observed in current mRNA vaccines ^33,34^. Although designing an mRNA vaccine with a relatively larger CUB-tRNA mismatch might reduce overall tRNA overconsumption, this does not necessarily mean that the translation efficiency of host endogenous genes will be unaffected. Our method may help predict which endogenous genes are more likely to be repressed by an mRNA vaccine, thereby assisting in the design of safer vaccines that minimize adverse impacts on important host genes.

## Materials and methods

### Ribo-seq data download and reanalysis

Raw sequencing reads from two datasets (Sample information listed in Table S1) were downloaded from NCBI BioProjects under accession numbers PRJNA1136820 (Ribo-Seq of pan-virally infected A549 cells) ^22^, PRJNA316125 (Ribo-Seq of wild type A549 cells) ^35^, and PRJNA1335396 (Ribo-Seq of wild type and GmR transgenic *E. coli*) ^16^. For these datasets, raw reads were processed by removing adapters using Cutadapt ^36^. The *E. coli* data were aligned to the reference genome (GCF_000005845.2) along with the corresponding *GmR* sequences (Table S2) using STAR ^37^. The A549 data were aligned to the reference genome (GCF_000001405.40) along with the customized viral sequences (Table S2) using STAR ^37^. Quality control and quantification of translation efficiency, which was defined as the transcripts per million (TPM) of ribosome-protected fragments (RPFs) per gene, were performed using RiboParser ^38^. Importantly, the translation efficiency is defined at the gene level (total RPFs), not at the level of individual mRNA molecules.

For any endogenous gene, the repression index is defined as the translation efficiency measured in the presence of the exogenous (viral or *GmR*) gene divided by that measured in the absence of the exogenous gene. A lower value of repression index means that the gene is more strongly inhibited by the exogenous ene.

### The genomes of human viruses download and reanalysis

In this study, the available genome sequences of viruses known to infect humans were used to evaluate the necessity of distinguishing among rare-tRNA shortage (RTS), abundant-tRNA shortage (ATS), and viral similarity repression (VSR). Viral sequences were downloaded from the NCBI Virus database ^39^. In detail, the viral sequences with human hosts were downloaded and extracted the corresponding virus species names (i.e., viruses confirmed to be capable of infecting humans). From this list, viruses that infect human gut microbes (e.g., bacteriophages) were removed, as these viruses target bacteria rather than human cells and are therefore not relevant to the virus–human tRNA competition examined in this study. Based on the remaining virus species names, the complete viral genome sequences were then retrieved from the NCBI Virus database, including both RefSeq and variant genomes. To ensure sequence integrity for subsequent analyses, only those genomes whose length was at least 80% of the length of the corresponding RefSeq genome for each virus species was retained. Notably, during this retrieval step, the host were not restricted to humans. Instead, for each virus species known to infect humans, the complete genome sequences obtained from any host (including humans, animals, and other sources) were downloaded. Finally, 60622 complete genome sequences of 524 human viruses were obtained. For SARS-CoV-2, due to the overwhelming number of available sequences, the sequences from non-human hosts (N = 561) and 1,000 randomly selected sequences from human host were included for the same analysis.

### CUB-tRNA mismatch of a given endogenous gene

In this work, the metric *D* were used to estimate the degree of mismatch between a gene’s codon usage and the cellular tRNA supply ^16^. Specifically, *D* is estimated in two ways: *D*_opt_ based on the codon usage bias (CUB) of highly expressed host genes, and *D*_tRNA_ based on the expression levels of tRNA genes. The overall *D* value for each endogenous gene (either *D_opt_* or *D_tRNA_*) is defined as the weighted geometric mean of all 18 *D*_i_ values, one for each amino acid with at least two synonymous codons. Each *D*_i_ is the Euclidean distance between the codon frequency vector of a given endogenous gene and the estimated tRNA supply of host for amino acid *i*, as follows:

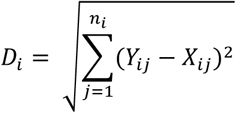

where *n_i_* is the number of synonymous codons for amino acid *i*, *Y_ij_* is the fraction of codon *j* among the synonymous codons for amino acid *i* in the endogenous gene (unweighted), and *X_ij_* is the estimated frequency of codon *j* in the host tRNA supply. For each amino acid, the sum of *X_ij_* (or *Y_ij_)* over all synonymous codons is 1.

The estimation of *X_ij_* differs between the two approaches: 1) For *D_opt_*, *X_ij_* is the expression-weighted frequency of codon *j* in the top 10% of highly expressed host genes:

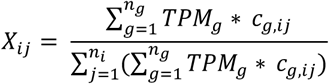

where *n_g_* is the total number of host genes that fall into the top 10% expression percentile, *g* is the index for a single host gene within that top 10% expression percentile, *TPM_g_* is the expression level of gene *g* measured in transcripts per million (TPM), *c_g,ij_* is the count of codon *j* (for amino acid *i*) in gene *g*, *n_i_* is the number of synonymous codons for amino acid *i*. Gene expression levels used to identify the top 10% of host genes were obtained from previous studies ^40,41^. 2) For *D_tRNA_*, *X_ij_* is estimated directly from tRNA expression levels. Specifically, *X_ij_* is defined as the normalized *W_j_* values (an absolute adaptiveness value) of codon *j* within its synonymous group of amino acid *i*. For each codon *j*, *W*_j_ was calculated analogous to the classic definition ^42^ but using tRNA gene expression levels (TPM) instead of copy numbers:

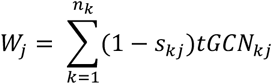

where *n_k_* is the number of tRNA isoacceptors that recognize codon *j*, tGCN_kj_ is tRNA gene expression levels (TPM) of tRNA *k* that recognizes codon *j*, *s*_kj_ is a selective constraint on the efficiency of the codon–anticodon coupling ^43,44^. The expression levels of each tRNA genes were extracted from previous publications ^45,46^.

Notably, the metric *D* above can also be applied to quantify the CUB-tRNA mismatch of a given endogenous gene in *E. coli*.

### A supplementary note of Cellular tRNA supply

Cellular tRNA supply ^16^ is a multifaceted concept that includes not only tRNA expression levels but also factors such as tRNA aminoacylation efficiency, the formation of the EFTu•GTP•aminoacylated-tRNA ternary complex, and codon–anticodon binding affinity (including wobble base pairing). Consequently, steady-state tRNA expression profiles alone may not accurately reflect the effective tRNA pool available for active translation. *D_opt_* is estimated from highly expressed human genes, which may not perfectly match the real tRNA pool because of the weak translational selection observed in metazoans ^47^. *D_tRNA_* is derived directly from total tRNA expression levels, which include non-translating tRNA pools.

This deviation from the actual tRNA pool may obscure weaker biological patterns, meaning that the patterns detected with these two imperfect metrics are likely to be robust underestimates of the true effects.

### Codon Usage Bias difference between a given endogenous gene and viruses

In this work, the metric *V* was used to estimate the difference in synonymous codon usage bias (CUB) between each endogenous gene and the viral genes. Only the 59 sense codons that belong to amino acids with at least two synonymous codons are considered (i.e., excluding stop codons, methionine, and tryptophan). The frequency of each codon is normalized within its synonymous group (i.e., for each amino acid, the frequencies of all its synonymous codons sum to 1). Formally,

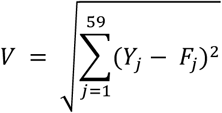

where *Y*_j_ is the frequency of codon *j* in a given endogenous gene (unweighted), and *F*_j_ is the expression-weighted (TPM of RPF) frequency of codon *j* aggregated across all viral genes. Specifically, *F*_j_ is calculated as:

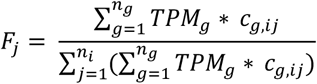

Here, *n_g_* is the total number of viral genes, *c_g,ij_* is the count of codon *j* (for amino acid *i*) in viral gene *g*, *n_i_* is the number of synonymous codons for amino acid *i*. *TPM_g_* is the expression level of gene *g* measured in TPM of RPF when such data are available ^22^; if expression data are unavailable, all *TPM_g_* are set to 1, resulting in an unweighted average of codon frequencies across viral genes. The resulting *V* is a single value for each endogenous gene, quantifying its overall CUB distance to the viral gene set. Notably, the same metric *V* can also be applied to quantify the CUB difference between a given endogenous gene of *E. coli* and the gentamicin resistance (*GmR*) gene.

## Declarations

## Ethics approval and consent to participate

Not applicable

## Consent for publication

Not applicable.

## Availability of data and materials

Custom R codes were used in data analysis and are available on GitHub (https://github.com/chenfengokha/viralRiboseq).

## Acknowledgements

Thanks are due to Jian-Rong Yang and Shanjun Deng for their comments on the manuscript. This work was supported by the National Key R&D Program of China (2022YFA1106700 to F.C.), the National Natural Science Foundation of China (32270681 to F.C.), and the Science and Technology Projects in Guangzhou (2025A04J5439 to F.C.)

## Authors’ contributions

F.C. conceived the idea, designed and supervised the study. F.C. acquired and analyzed data. F.C. wrote the paper.

---

**Fig. S1.**
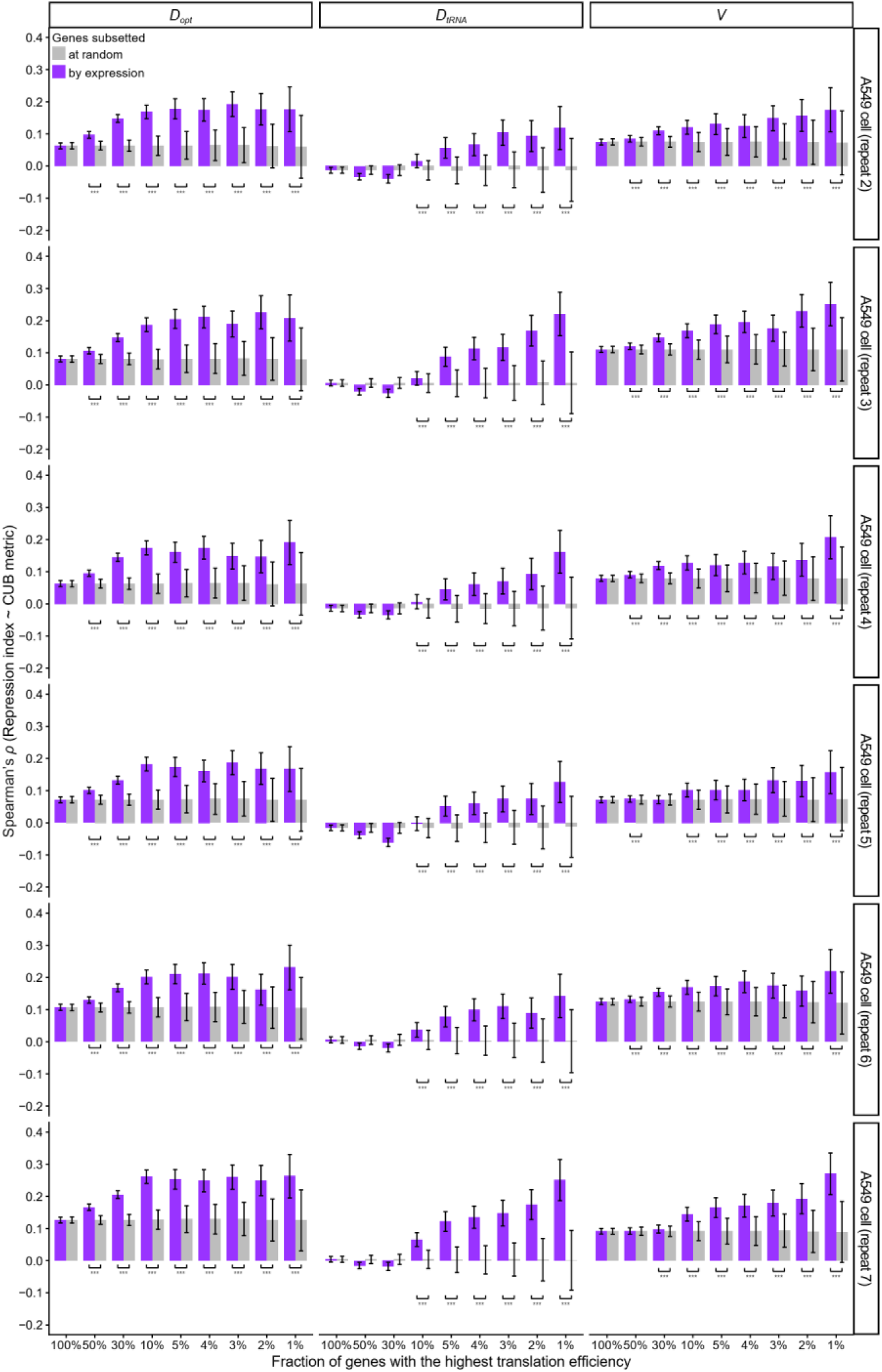
Additional biological replicates reproduce the correlations seen in Fig. 2. For *D_opt_* (left), *D_tRNA_* (middle), and *V* (right), Spearman correlations between repression index and each metric were calculated across translation efficiency thresholds. The positive correlations strengthen with increasing thresholds, and random downsampling controls (gray bars) rule out gene-number effects, confirming the robustness of the main findings.

**Fig. S2.**
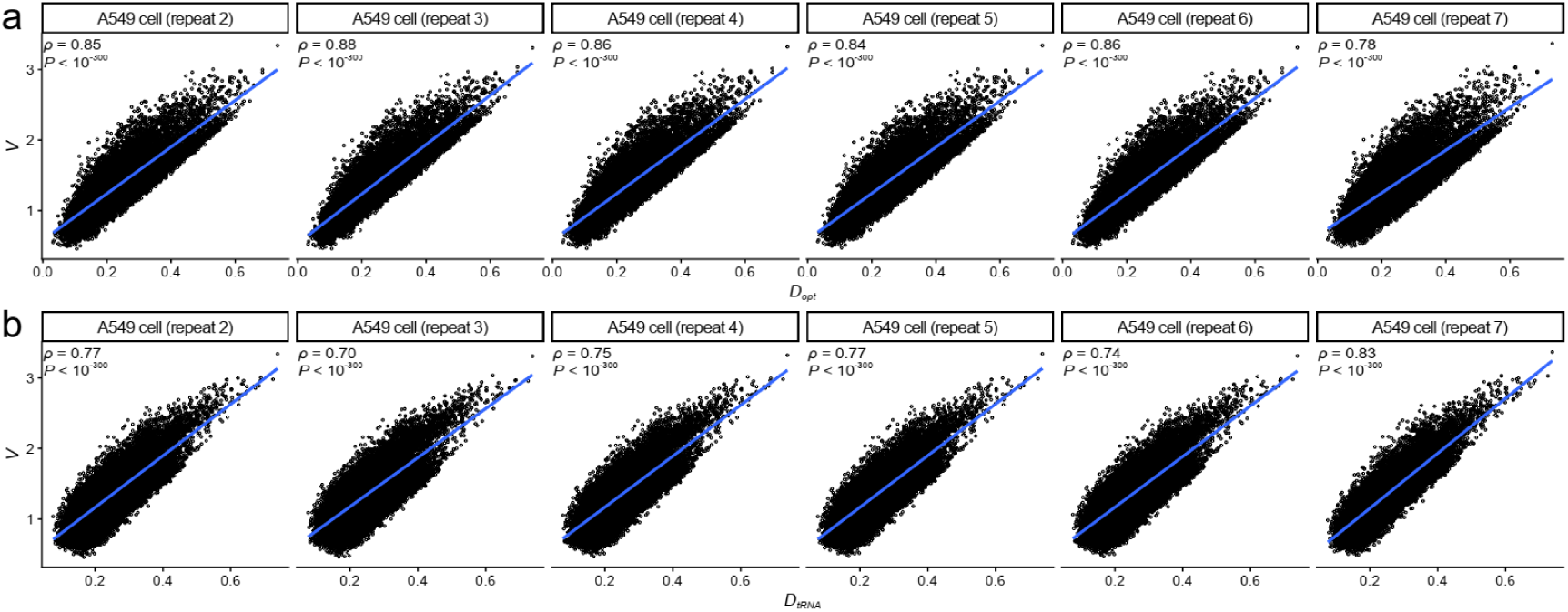
Additional replicates reproduce the *V*-*D_opt_* and *V*-*D_tRNA_* correlations seen in Fig. 3d-e. For each of the biological replicates, Spearman correlation analysis shows that *V* is significantly positively correlated with *D_opt_* (a) and with *D_tRNA_* (b).

**Fig. S3.**
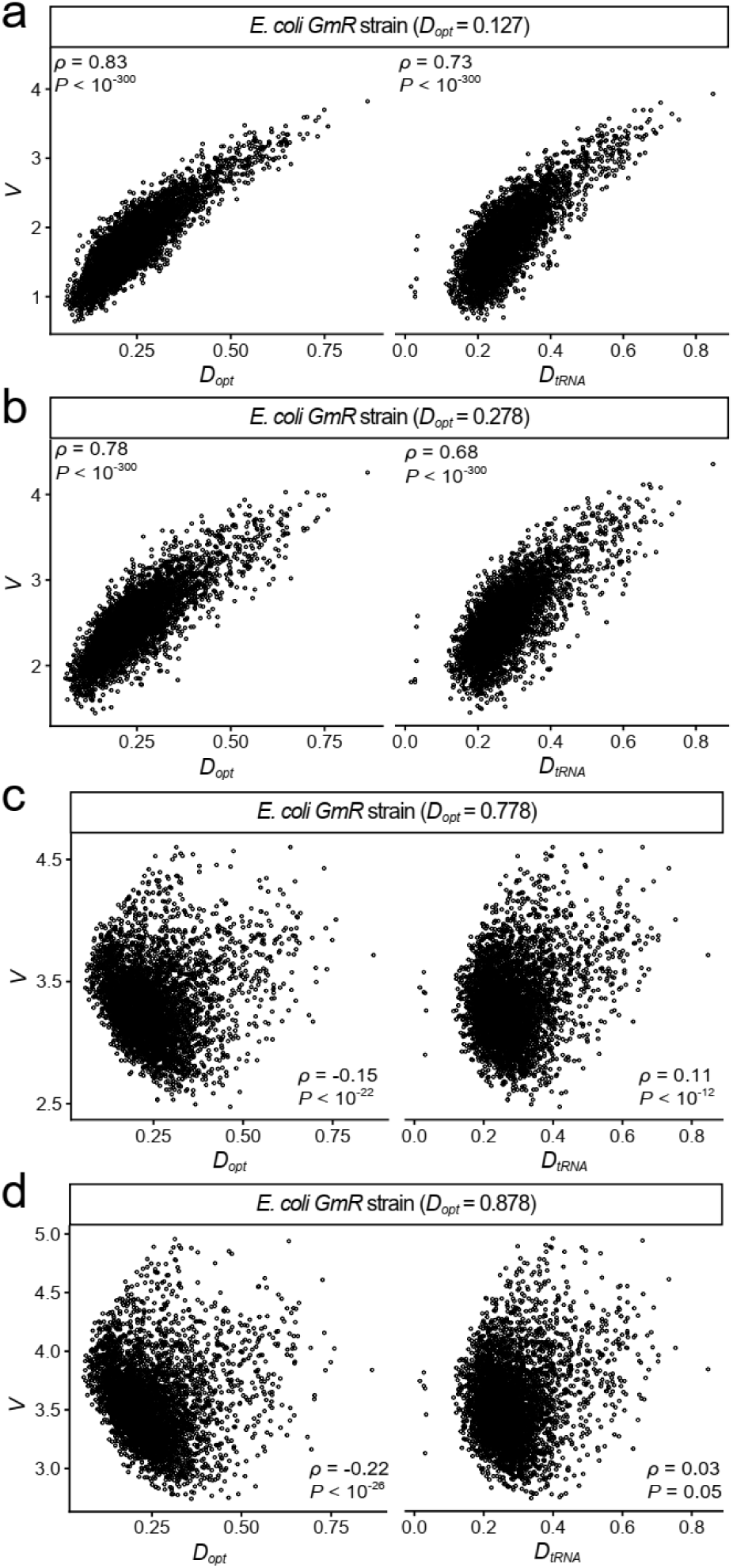
Spearman rank correlations between *V* and CUB-tRNA mismatch (Dopt, DtRNA) of endogenous genes across the four *GmR*-expressing *E. coli* strains. (a-b) *GmR* strains with small CUB-tRNA mismatch (*D_opt_* = 0.127 and 0.278). (c-d) *GmR* strains with large CUB-tRNA mismatch (*D_opt_* = 0.778 and 0.878). The two-tailed Spearman rank correlation coefficients (two-tailed) and *P* values were shown.

**Fig. S4.**
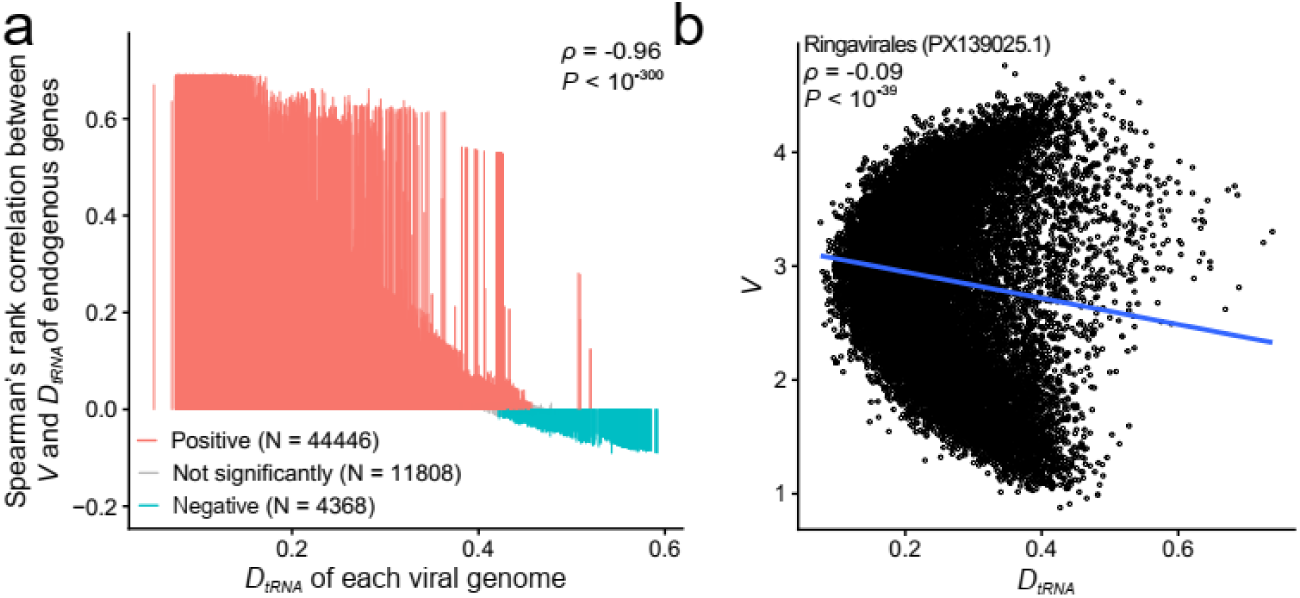
The same analysis as Fig. 5 using another alternative CUB-tRNA mismatch metric *D_tRNA_*. (a) Lollipop chart analogous to Fig. 5a, but using *D_tRNA_* instead of *D_opt_*. (b) Representative example of a virus showing a negative correlation between endogenous *V* and *D_tRNA_*. The virus name and corresponding GenBank accession number are indicated in the panel. Each point represents an endogenous gene.

**Table S1.**
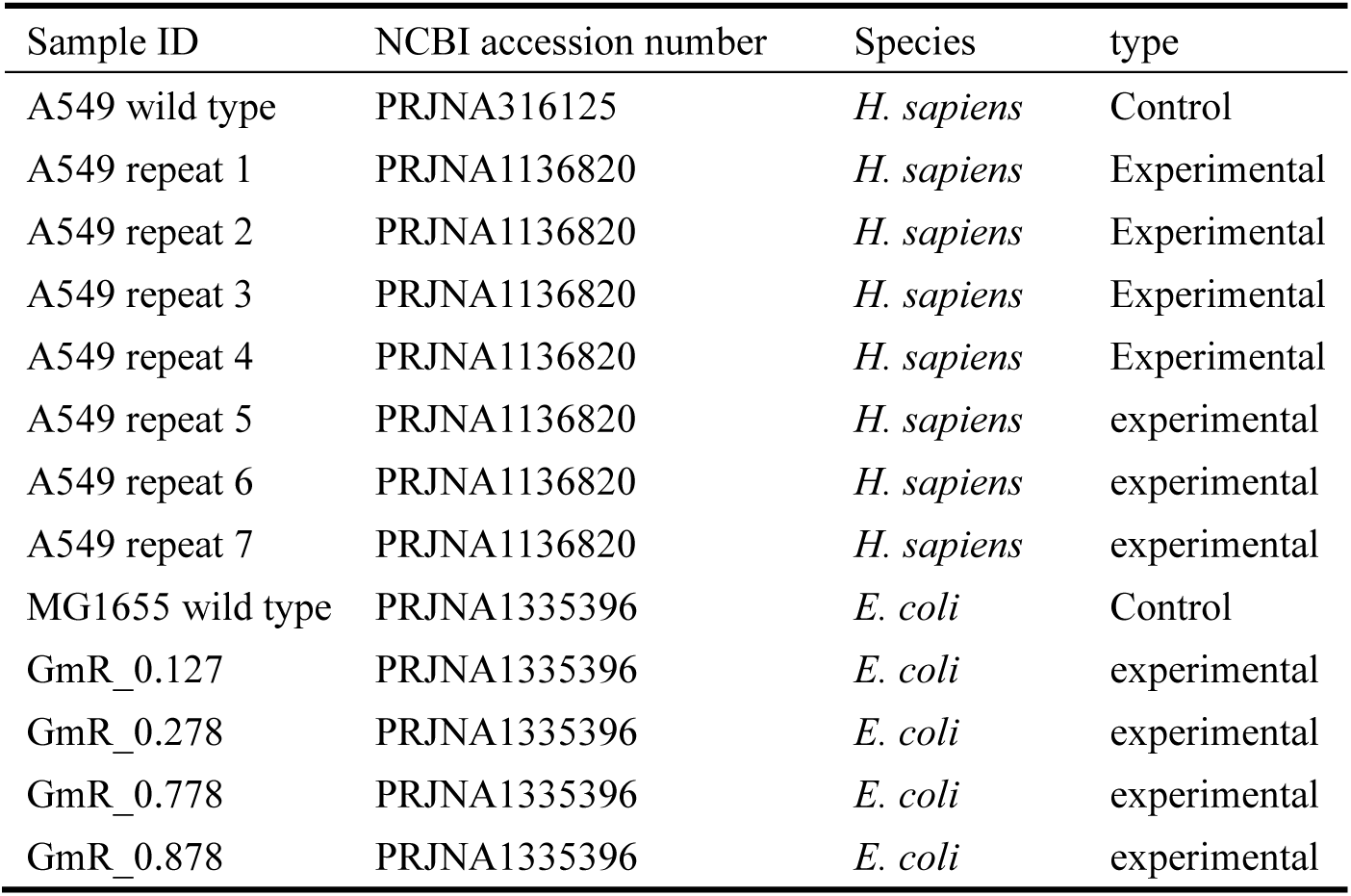
Sample information.

| Sample ID | NCBI accession number | Species | type |
| --- | --- | --- | --- |
| A549 wild type | PRJNA316125 | *H. sapiens* | Control |
| A549 repeat 1 | PRJNA1136820 | *H. sapiens* | Experimental |
| A549 repeat 2 | PRJNA1136820 | *H. sapiens* | Experimental |
| A549 repeat 3 | PRJNA1136820 | *H. sapiens* | Experimental |
| A549 repeat 4 | PRJNA1136820 | *H. sapiens* | Experimental |
| A549 repeat 5 | PRJNA1136820 | *H. sapiens* | experimental |
| A549 repeat 6 | PRJNA1136820 | *H. sapiens* | experimental |
| A549 repeat 7 | PRJNA1136820 | *H. sapiens* | experimental |
| MG1655 wild type | PRJNA1335396 | *E. coli* | Control |
| GmR_0.127 | PRJNA1335396 | *E. coli* | experimental |
| GmR_0.278 | PRJNA1335396 | *E. coli* | experimental |
| GmR_0.778 | PRJNA1335396 | *E. coli* | experimental |
| GmR_0.878 | PRJNA1335396 | *E. coli* | experimental |

**Table S2.**
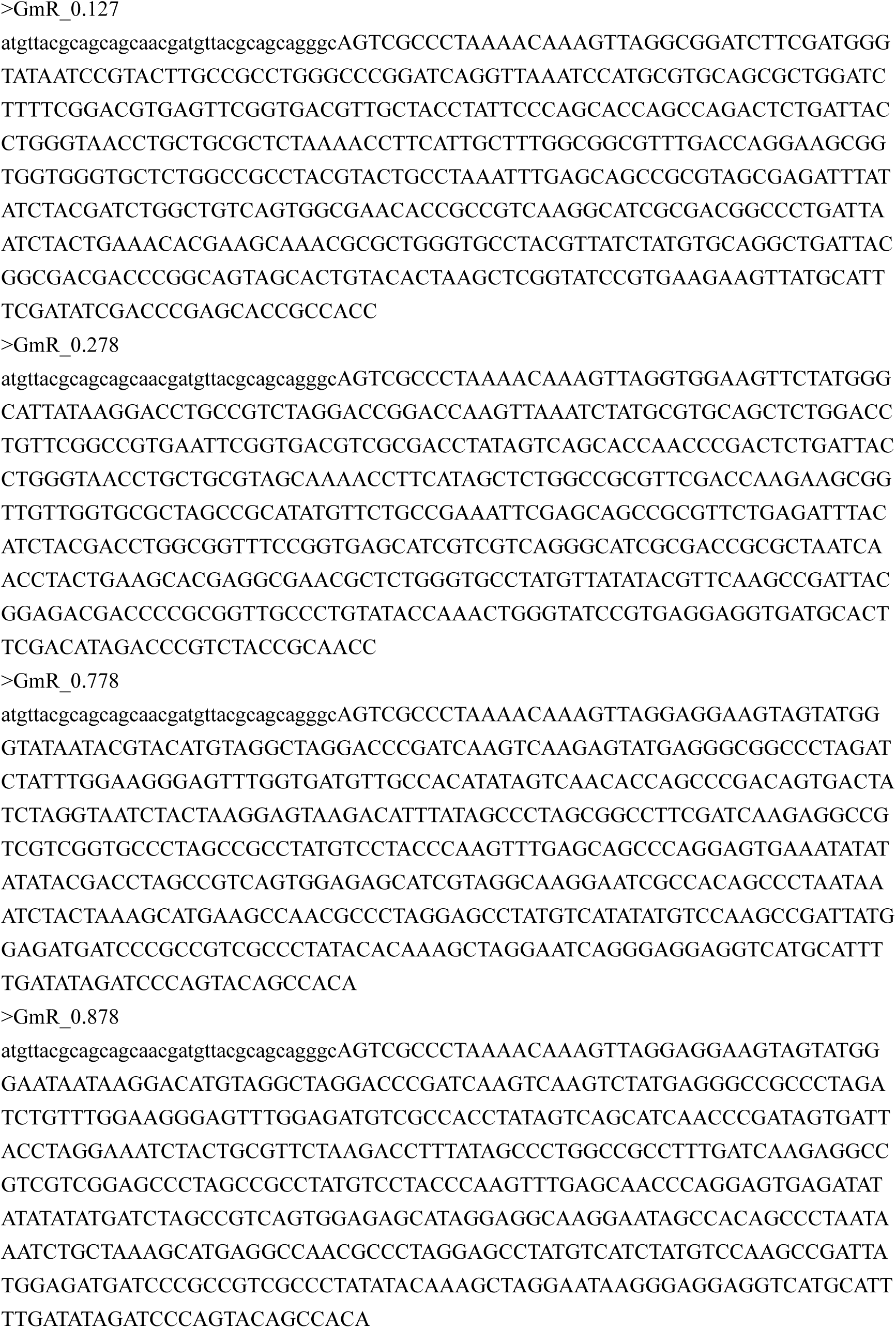
*GmR* (N = 4) and viral sequences (N = 3532; too many to show).

**Table S3.**
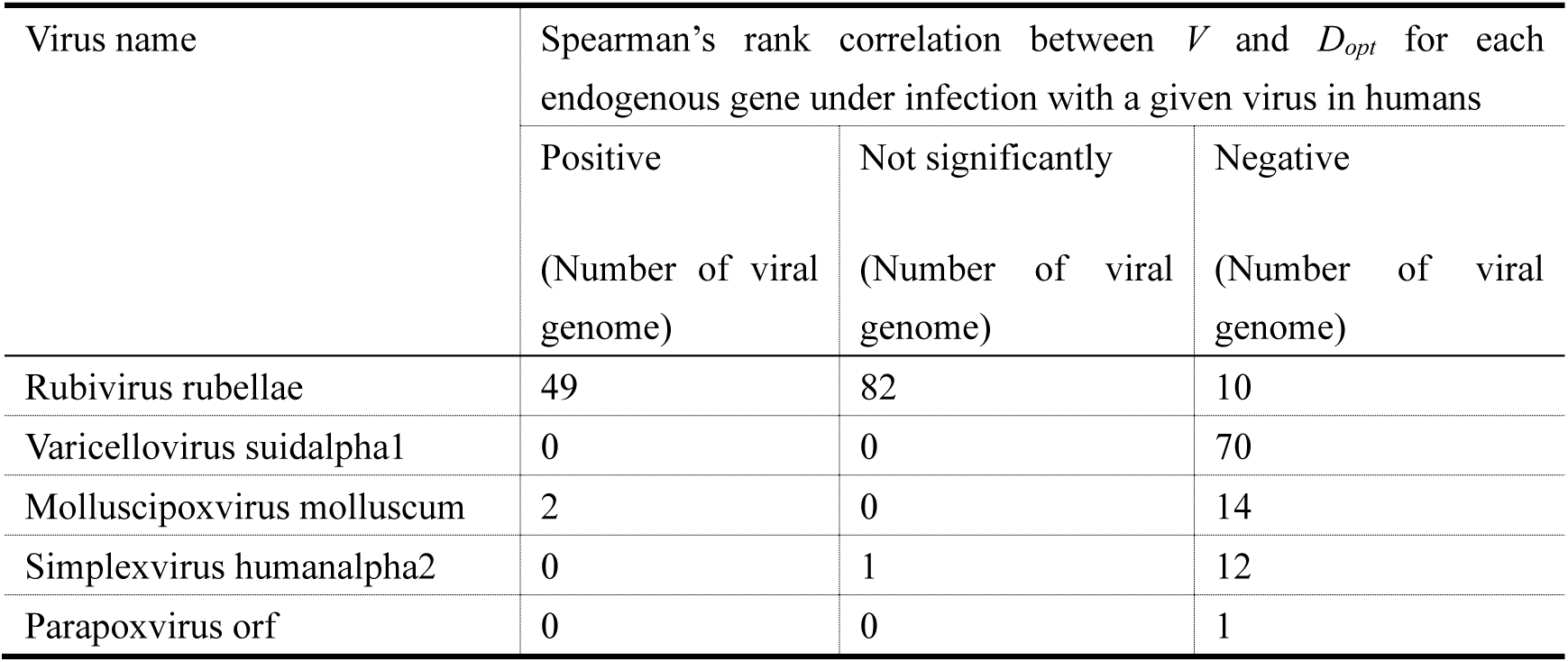
Viruses show non-positive correlation between endogenous *V* and *D_opt_*.

**Table S4.**
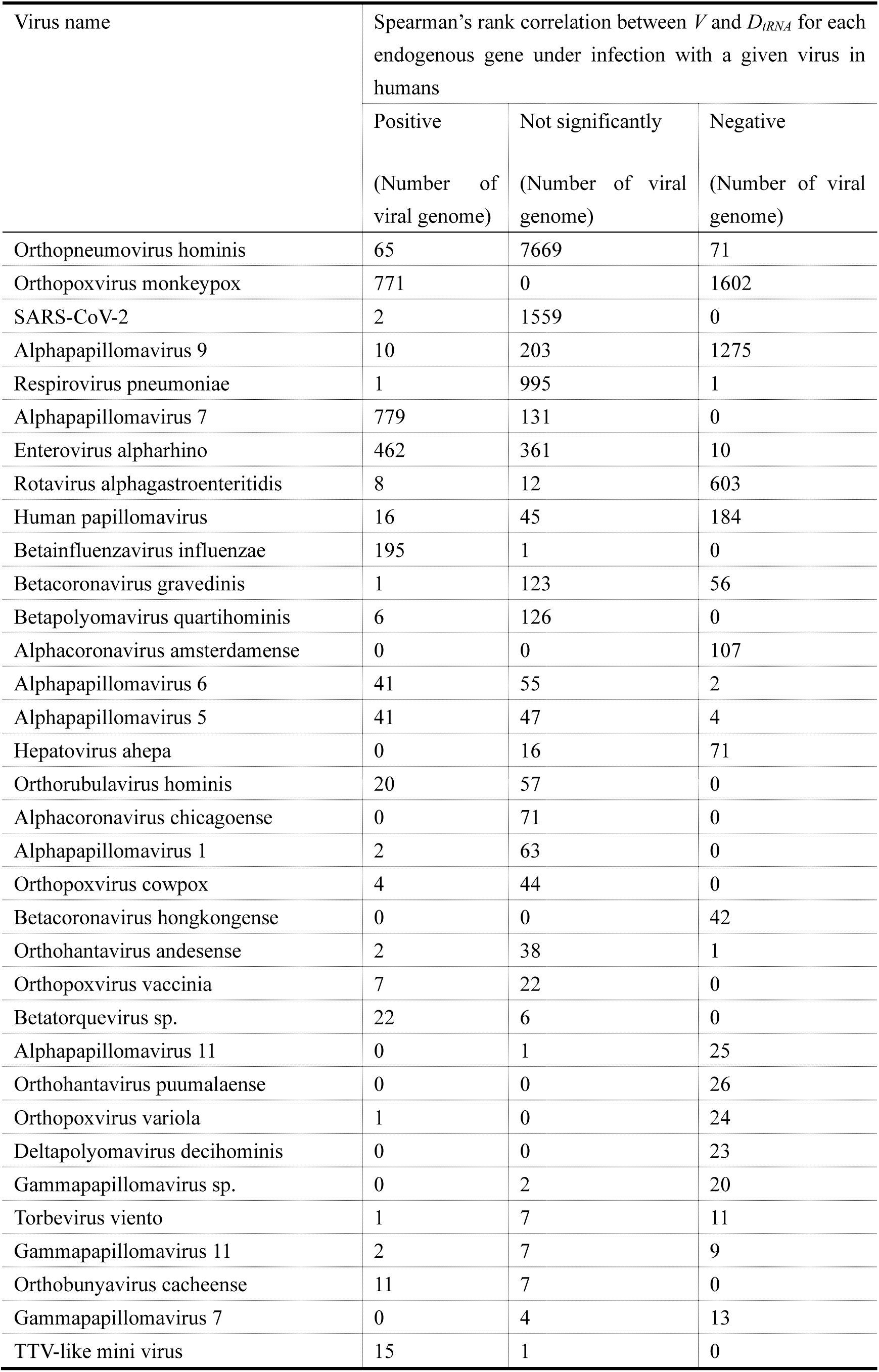
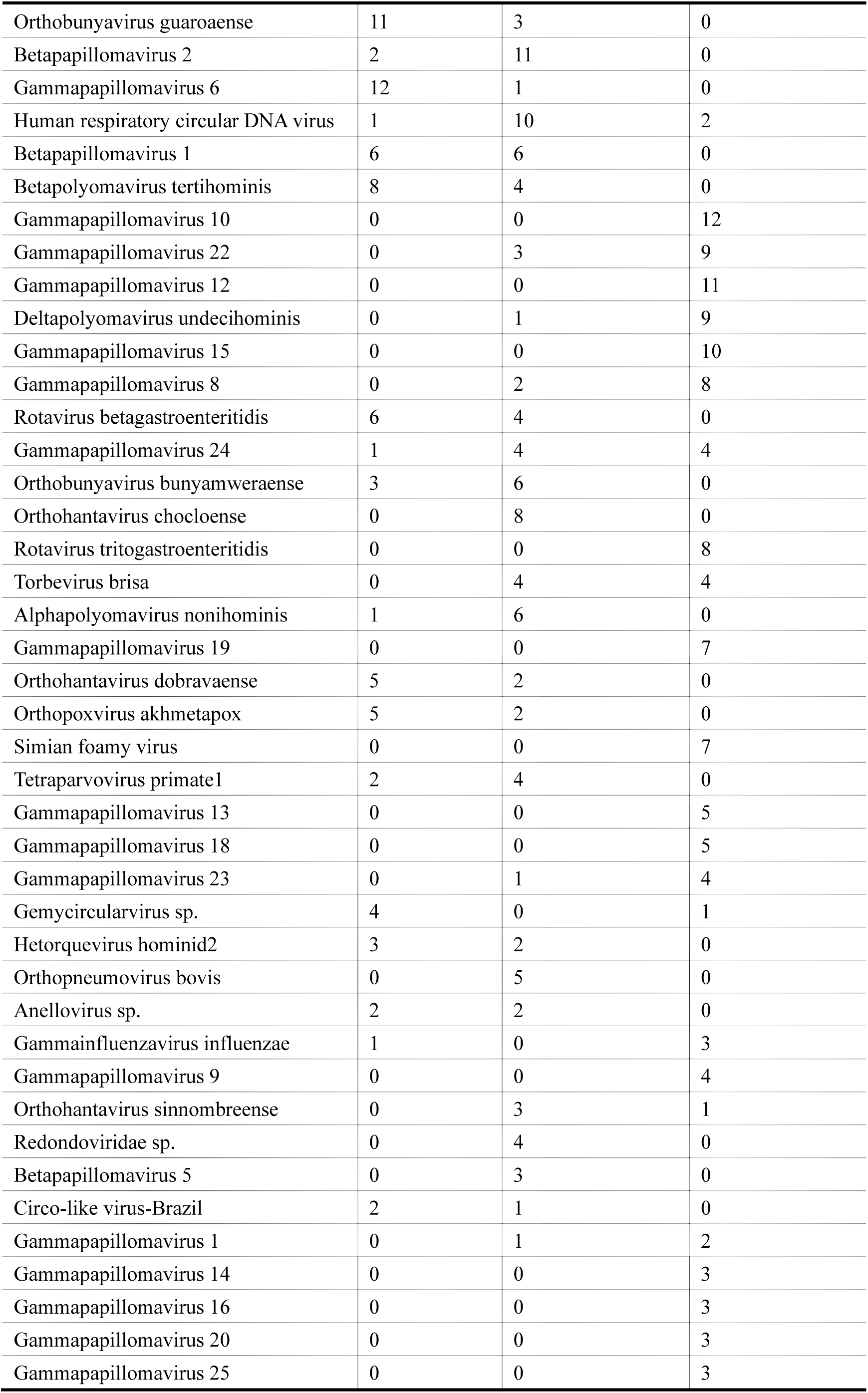
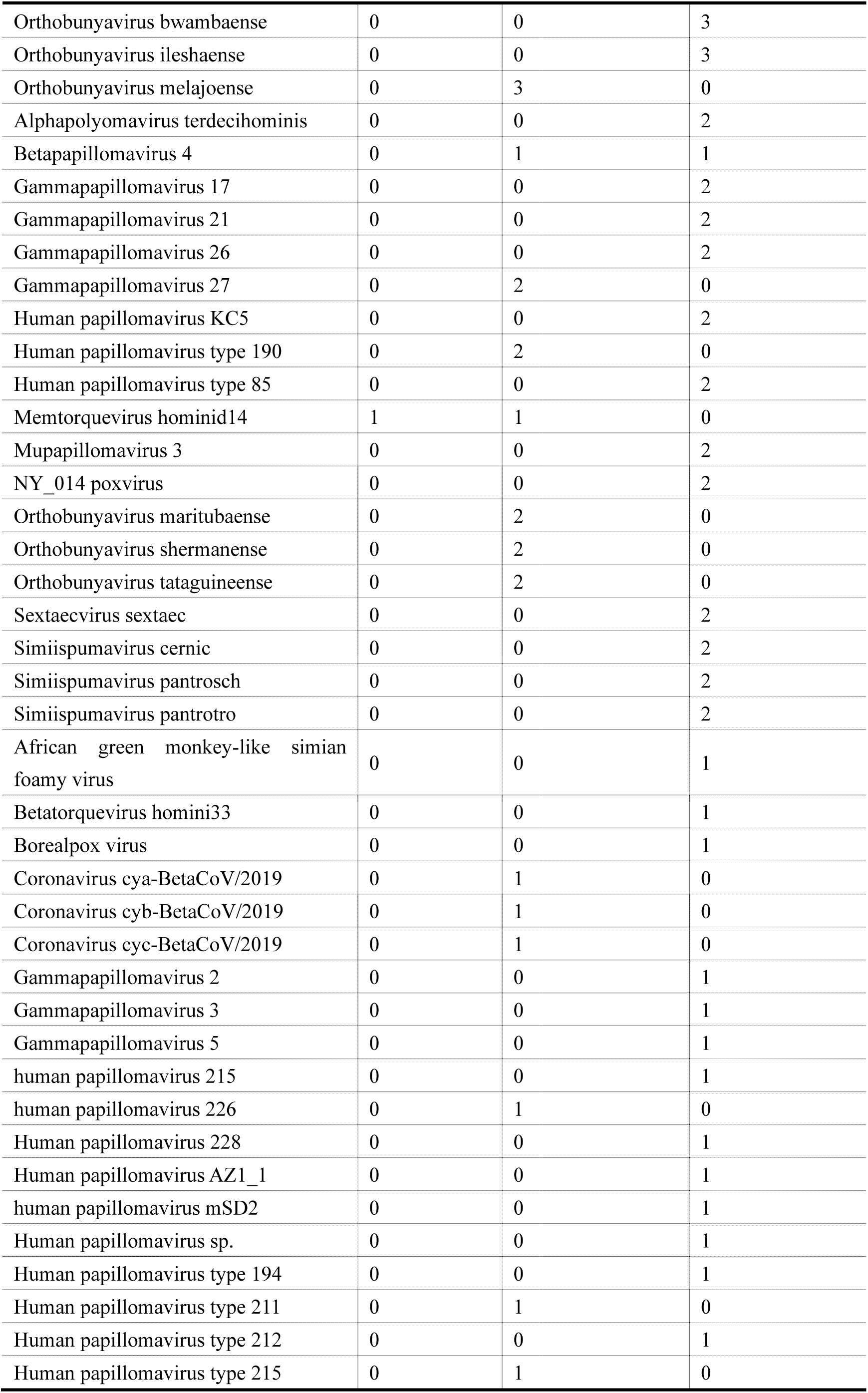
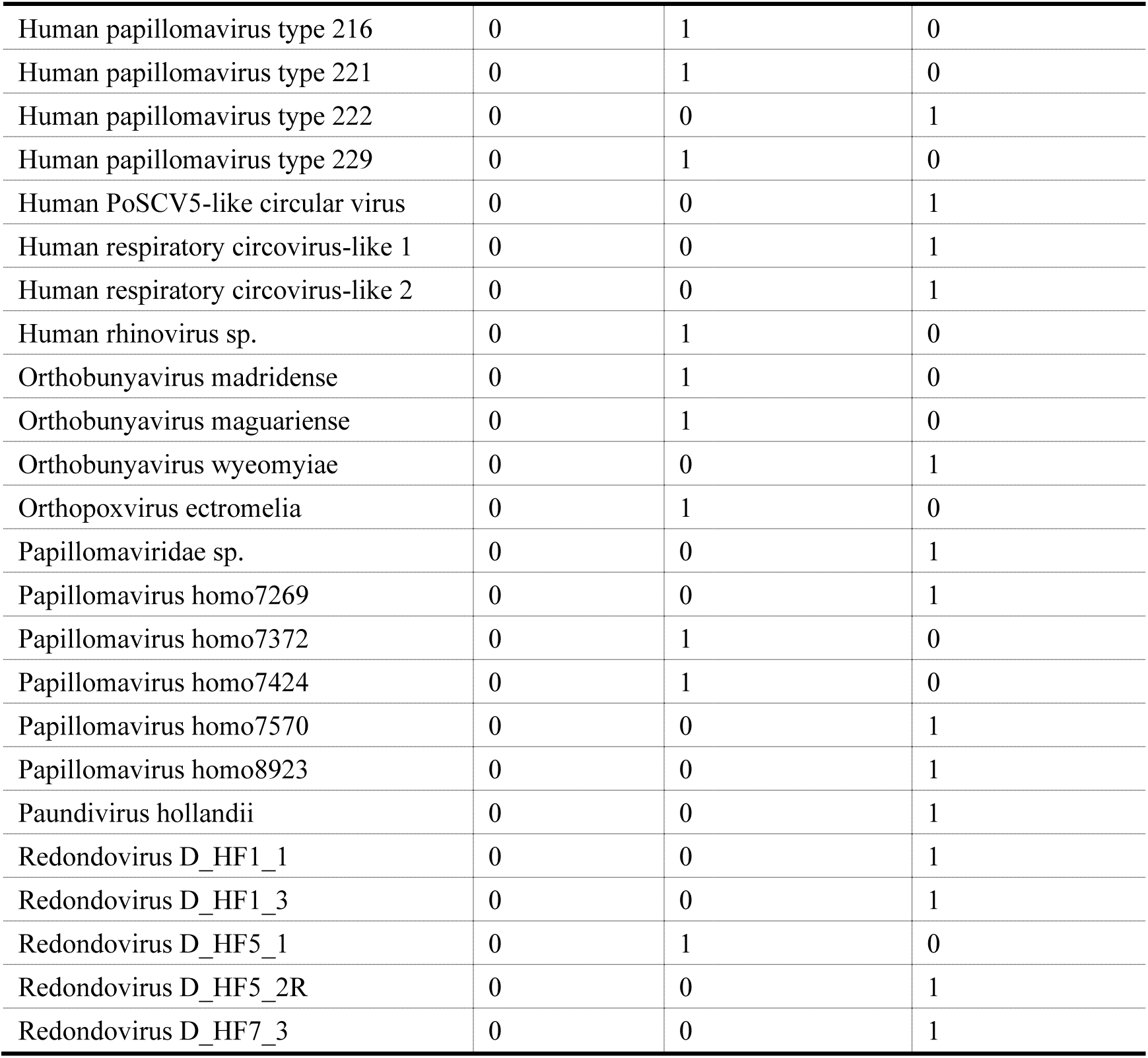
Viruses show non-positive correlation between endogenous *V* and *D_tRNA_*.

